# Alpha-Interferons and CD40: complementary pathways revealed by mGEDO in distinct transcripto-methylomic profiles of patients with Sjögren’s disease

**DOI:** 10.64898/2026.11.25.768874

**Authors:** Clément Bézier, Jakez Rolland, Ronan Boutin, Divi Cornec, Sophie Hillion, Christelle Le Dantec, Eleonore Bettacchioli, Xavier Mariette, Jacques-Eric Gottenberg, PRECISESADS Clinical Consortium, Nathan Foulquier, Valérie Devauchelle-Pensec

## Abstract

**Objectives:** Sjögren’s disease (SjD) exhibits substantial clinical and molecular heterogeneity, challenging patient stratification and targeted therapies. We aimed to develop multi-omic signatures of interferon-alpha (IFN) and CD40 pathways using mGEDO, and characterize their activity across SjD subgroups and healthy controls.

**Methods:** We developed mGEDO to integrate whole-blood transcriptomic and DNA methylation data from PRECISESADS using BloodGen3 modules, generating continuous two-dimensional patient representations. IFN and CD40 signatures were derived through automated gene selection and quantified using mGEDO Transition Scores. Associations with clinical and biological phenotypes were assessed in PRECISESADS and independently investigated in ASSESS.

**Results:** Among 702 PRECISESADS participants, mGEDO revealed a continuous SjD–control molecular landscape with substantial heterogeneity beyond discrete clustering. IFN and CD40 signatures exhibited distinct, partially overlapping patterns. IFN was predominantly associated with biological disease activity, whereas CD40 provided complementary information, particularly in IFN-negative patients. Combined IFN/CD40 stratification identified biologically progressive subgroups, with double-positive patients exhibiting the greatest immune dysregulation. In 371 ASSESS patients, *CD40* and *CD40LG* expression associations with biological phenotypes were confirmed. Both genes were associated with circulating immune-cell composition.

**Conclusions:** IFN and CD40 signatures reflect complementary, partially uncoupled innate and adaptive immune programs, potentially improving molecular stratification and precision therapeutic targeting in SjD. Whole-blood *CD40LG* expression may reflect immune-cell redistribution and serve as a potential marker of immune-cell migration.

## 1. Introduction

Sjögren’s disease (SjD) is a systemic autoimmune disease characterized by lymphoid infiltration and tissue damage in the salivary and lacrimal glands, leading to severe and handicapping global dryness (Teruel et al., 2021). Clinically, SjD patients present with great heterogeneity, highlighted in clinical and biological manifestations (Nguyen et al., 2024), but also in the molecular profiles, both in the blood and in the glands (Soret et al., 2021). This heterogeneity in the clinical and serological manifestations leads to classification criteria based on a multi-level score, using a differential diagnosis exclusion, factual evaluation of dryness, and SSA autoantibodies (Shiboski et al., 2017). Landscape of autoantibodies is mainly based on TROVE2 / Ro60, TRIM21 / Ro52 (specifically related to Interferon signature and severity of the disease (Bettacchioli et al., 2024)), SSB and rheumatoid factors. But not all patients reaching classification criteria present anti-SSA autoantibodies. Systemic manifestations also occurred in 30 to 40 percent of SjD patients with different manifestations such as skin, joints, neurologic, lung, or kidney involvement (Mariette and Criswell, 2018). This heterogeneity has a real impact on the efficacy of treatments (Soret et al., 2021; Chevet et al., 2025). So, experts considered that distinct molecular signatures and pathophysiology of SjD could lead to the choice of personalized treatment, depending on the biological pathways involved (Barturen et al., 2018; Chevet et al., 2025).

For years, the type-I interferon signature has been identified as a characteristic of the disease and is associated with anti-SSA autoantibody (Gottenberg et al., 2006; Teruel et al., 2021; Soret et al., 2021; Trutschel et al., 2022), NF-kB signaling pathways, and T cell activation. But not all patients revealed interferon pathway activation, with some authors suggesting a role of different molecular or immunological pathways (Del Papa et al., 2021).

Recently, efficacy of therapies targeting CD40 or CD40L (Iscalimab, Dazodalibep (Fisher et al., 2020, 2024; St. Clair et al., 2024)) highlighted the role of the CD40-CD40L pathway in SjD, but this pathway has not yet been studied at the patient level. Initially described as a B cell growth factor (Clark and Ledbetter, 1986), later named CD40 (Paulie et al., 1989), the CD40-CD40L (or CD154) axis seemed very attractive in the field of organ transplantation and autoimmune diseases (such as systemic lupus erythematosus, rheumatoid arthritis, and SjD), with CD40L leading to the activation of CD40-expressing cells (Pucino et al., 2020). CD40-CD40L axis is related to follicular T cells (Tfh) - B cells cooperation and is essential for the formation of germinal centers, immunoglobulin class switching, and B cell activation (Crotty, 2011). The protein encoded by the *CD40LG* gene is expressed at the surface of T cells and regulates B cell function through the interaction with CD40 expressed at the B cell surface (Liu et al., 2022). Since 2019, the rational for targeting the CD40-CD40L axis has been developed in autoimmune diseases (Karnell et al., 2019; Pucino et al., 2020) and in SjD particularly (Pucino et al., 2020). CD40L antagonist (St. Clair et al., 2024) and anti-CD40 antibody (Fisher et al., 2020, 2024) are currently under evaluation.

Identifying robust SjD subgroups or endotypes remains challenging. Clinical-symptom-based classifications (McCoy et al., 2022) lead to unclear molecular profiles, even associated with biology (Nguyen et al., 2024), and molecular clustering (Soret et al., 2021; Trutschel et al., 2022; Barturen et al., 2021) can lead to complex clinical characterization. In general, different clustering approaches can lead to non-consensus classification across studies. It exists a substantial overlap between SjD and healthy controls and among SjD themselves in stratification studies (Soret et al., 2021). Emerging work axis pose the hypothesis of several disease stages: initiation, preclinical and asymptomatic, overt SjD and chronic disease, underlining that these different autoimmune processes are incompletely known (Wang et al., 2021). For these reasons, discrete clusters may categorize the information without accurately capturing the continuum of SjD and CTRL patients and the transitions between SjD subgroups. We suggest that SjD could be a continuum between healthy controls and most severe patients, and that different subgroups of SjD could be complexly connected in different regions with transitions between each other, suggesting the use of a continuous landscape of SjD rather than discrete categories to better understand all its heterogeneity. We thus introduce mGEDO, the multi-omic extension of our previous GEDO algorithm: a gene set analysis method based on a transition score computing the position of individuals on the continuum between healthy controls and diseased patients (Bézier et al., 2026).

PRECISESADS is a large European multicenter cohort aimed at identifying molecular signatures to stratify systemic autoimmune diseases using a wide range of omic analyses (Barturen et al., 2018). This gave us the opportunity to evaluate large-scale whole-blood RNA sequencing and CpG methylation in patients with SjD and healthy controls. Using the mGEDO multi-omic extension, our objective was to propose a change of paradigm: representing SjD patients in a continuous landscape by the construction of two-dimensional visualizations of patients, rather than clustering patients into categories. This approach makes it possible to visualize continuums, gradual transitions, and pathways activation between different groups of patients in the PRECISESADS cohort.

Specific signatures were designed and computed for alpha-Interferons (IFN) and CD40, integrating both transcriptomic and methylomic data.

## 2. Methods

### 2.1. Patient cohorts

#### Exploratory dataset: PRECISESADS cohort

The PRECIS-ESADS cohort is a European multicenter study designed to reclassify Systemic Autoimmune Diseases -Systemic Lupus Erythematosus, Systemic Sclerosis, SjD, Rheumatoid Arthritis, Primary Antiphospholipid Syndrome, and Mixed and Undefined Connective Tissue Disease-on the basis of a large and diverse set of molecular data, including genomic, transcriptomic, epigenomic, metabolomic, and clinical information, with the aim of identifying SjD endotypes and novel molecular biomarkers, thereby improving the efficacy of innovative treatments targeting specific endotypes (Barturen et al., 2018) (ClinicalTrials.gov: NCT02890121). Healthy controls (CTRL) are defined as individuals with no history or current cases of autoimmune disease, chronic medication use, allergic or infectious conditions, thyroid disease, or any other condition that could alter the cellular composition of the blood (Barturen et al., 2021). We used the cross-sectional cohort of 2014 individuals with bulk RNA-seq data, including 258 SjD patients and 444 CTRL individuals. Among these 702 individuals, only 335 (144 SjD and 191 CTRL) had CpG methylation data (M-values) available (see Supplementary Notes for patient involvement statement, data preprocessing, and batch correction).

#### Validation dataset: ASSESS cohort

ASSESS cohort is a French national multi-center prospective cohort of patients with SjD (ClinicalTrials.gov: NCT03040583). Since 2006, patients have been followed for twenty years for the purpose of detecting new markers to predict the appearance of lymphoma subsequent to SjD (Duret et al., 2023). At baseline, 395 patients with SjD (fulfilling American-European Consensus Criteria (AECG) between 2006 and 2009) were included, with a large amount of clinical information and whole-blood microarray transcriptomic profiling for 371 of them. Preprocessing of ASSESS microarray data is detailed in the supplementary notes of the work of (Trutschel et al., 2022).

### 2.2. A mapping of SjD and CTRL with mGEDO

Our first objective was to identify different SjD subgroups via unsupervised analyses, and to identify pathways responsible for their differences with the CTRL group. First, singleomic analyses were performed on RNA-seq and methylomic data with GEDO (Bézier et al., 2026) (Supplementary Notes). Then, multi-omic integration was carried out with mGEDO, which reduces the dimensionality of RNA-seq and methylomic data by integrating them at the gene-set level. Both single- and multi-omic analyses were run using the 382 BloodGen3 gene sets (Altman et al., 2021).

Method for multi-omic integration: for each gene set, (1) corresponding genes were selected from RNA-seq data, and data for each CpG were selected if they correspond to regions of interest of these genes. (2) RNA-seq and CpG subsets were integrated using the NEMO similarity network fusion method (Rappoport and Shamir, 2019), allowing the integration of individuals with missing methylomic data. The output of NEMO is an integrated similarity network, with each node representing an individual (SjD or CTRL) and edges representing their multi-omic similarity. (3) After preprocessing, the resulting similarity network was used as input to compute a Transition Score (TS) based on the GEDO method (Bézier et al., 2026). TS is computed for each point *x* as its relative position to the CTRL and SjD groups in the similarity network. TS is a supervised score with: TS(*x*) ≈ 0: *x* is much closer, on average, to the CTRL group than to the disease group (SjD). TS(*x*) ≈ 0.5: *x* is approximately equidistant from both the CTRL and disease (SjD) group. Such individuals may lie in a transition zone and do not clearly resemble either group. TS(*x*) ≈ 1: *x* is much closer, on average, to the disease group (SjD) than to the CTRL group. Details of the mGEDO method and parameters used for this study are provided in the Supplementary Notes. This leads to the construction of a new gene-module matrix including SjD and CTRL and gene-module scores, which have been projected in two dimensions for direct visualization with UMAP and PHATE (two complementary visualization methods; see Supplementary Notes for hyperparameters chosen). These projections are called “mapping”. While UMAP (McInnes et al., 2018) reveals compact, clearly distinct clusters, PHATE (Moon et al., 2019) helped us understand the over-all structure of the transitions or trajectories between clusters. Correlations between each annotated gene module in the module matrix and PHATE1/PHATE2 and UMAP1/UMAP2 axes were computed in order to visualize which biological functions (among BloBloodGen3 modules) were the most important in the construction of the mapping.

### 2.3. IFN and CD40 signatures in SjD patients and CTRL

The objective was to derive multi-omic signatures representative of the IFN and CD40 pathways by integrating gene expression and CpG methylation data. The well-characterized IFN pathway (Del Papa et al., 2021; Teruel et al., 2021; Trutschel et al., 2022) was used as a reference to evaluate the CD40 signature in comparison.

To select gene sets, gene candidates were selected according to their representativeness and specificity to pathways under study. For this, we developed an automatic gene set selection tool exploring the GSEA database using keywords: “IFNA” and “INTERFERON_ALPHA” for IFN, and “CD40” for CD40 in our case (Supplementary Notes).

The resulting gene sets were used to extract RNA-seq and CpG methylation data. Three complementary scoring strategies were then applied. (1) Mean z-scores relative to CTRL quantified global expression or methylation changes. (2) Regularized Canonical Correlation Analysis (rCCA, mixOmics (Rohart et al., 2017)) was trained on the 335 individuals with paired RNA-seq and methylation data to derive one RNA and one CpG component maximizing cross-omic correlation. While MZS captures global regulation and rCCA extracts the main correlated sources of variation, neither method specifically targets disease-related alterations. (3) Therefore, supervised TS were computed on RNA-seq and DNA methylation separately, and then integrating the two, providing pathological axes that quantify each individual’s similarity to CTRL or SjD. The resulting integrated scores, TS_XY_IFN_ and TS_XY_CD40_, are hereafter referred to as the IFN and CD40 signatures.

Even if our hypothesis of continuums remains, to focus on the complementarity between IFN and CD40 signature and compute comparative statistics, individuals were classified as IFN- or CD40-positive using the 95th percentile of TS_XY scores in the CTRL group as the threshold. Combining both binary classifications defined four CD40/IFN groups: CD40+/IFN+, CD40+/IFN−, CD40 −/IFN+, and CD40 −/IFN−.

In parallel, the *TNFSF13B* gene expressing BAFF and the *TNFRSF13C* gene expressing BAFF-R were analyzed to make a parallel with BAFF pathway and its markers on B and T cells.

For validation of the signatures, gene-sets specificity was assessed by overlap analysis, and the multi-omic TS signatures were compared with published signatures and gene sets (Supplementary Notes).

As transcriptomic analyses are performed on whole blood, gene counts are intrinsically sensitive to cell composition changes if they are expressed by particular cell types. To understand the contribution of cell composition in *CD40* and *CD40LG* global expression, we used linear models to predict *CD40* and *CD40LG* with two panels of cells, one general panel, and one B cell-specific panel. P-values were corrected with the FDR method (Supplementary Notes).

### 2.4. Analysis of clinical and biological profiles in PRECIS-ESADS and ASSESS cohorts

To test whether SjD patients had different clinical profiles according to CD40 and IFN signatures, 91 features in the xzPRECISESADS cohort, including medical history, comorbidities, clinical examinations, biological parameters, treatments, symptoms, and autoantibodies, were tested against CD40-IFN class (Supplementary Notes). Statistics were computed only in SjD patients. To confirm the CD40 signature and its association with clinical profiles of patients, we also tested for associations between CD40 genes in our gene set and the huge number of clinical and biological characteristics at baseline in the ASSESS cohort, as a validation dataset. In addition, we analyzed ESSDAI and ESSPRI scores at different times (in months) after recruitment. Among all features available, we selected a subset of 225 characteristics of interest for SjD profiling (Supplementary notes). All p-values were corrected for multiple testing.

Figure 1 summarizes all the analyzes on mapping, signatures, and analysis in the ASSESS cohort.

**Figure 1.**
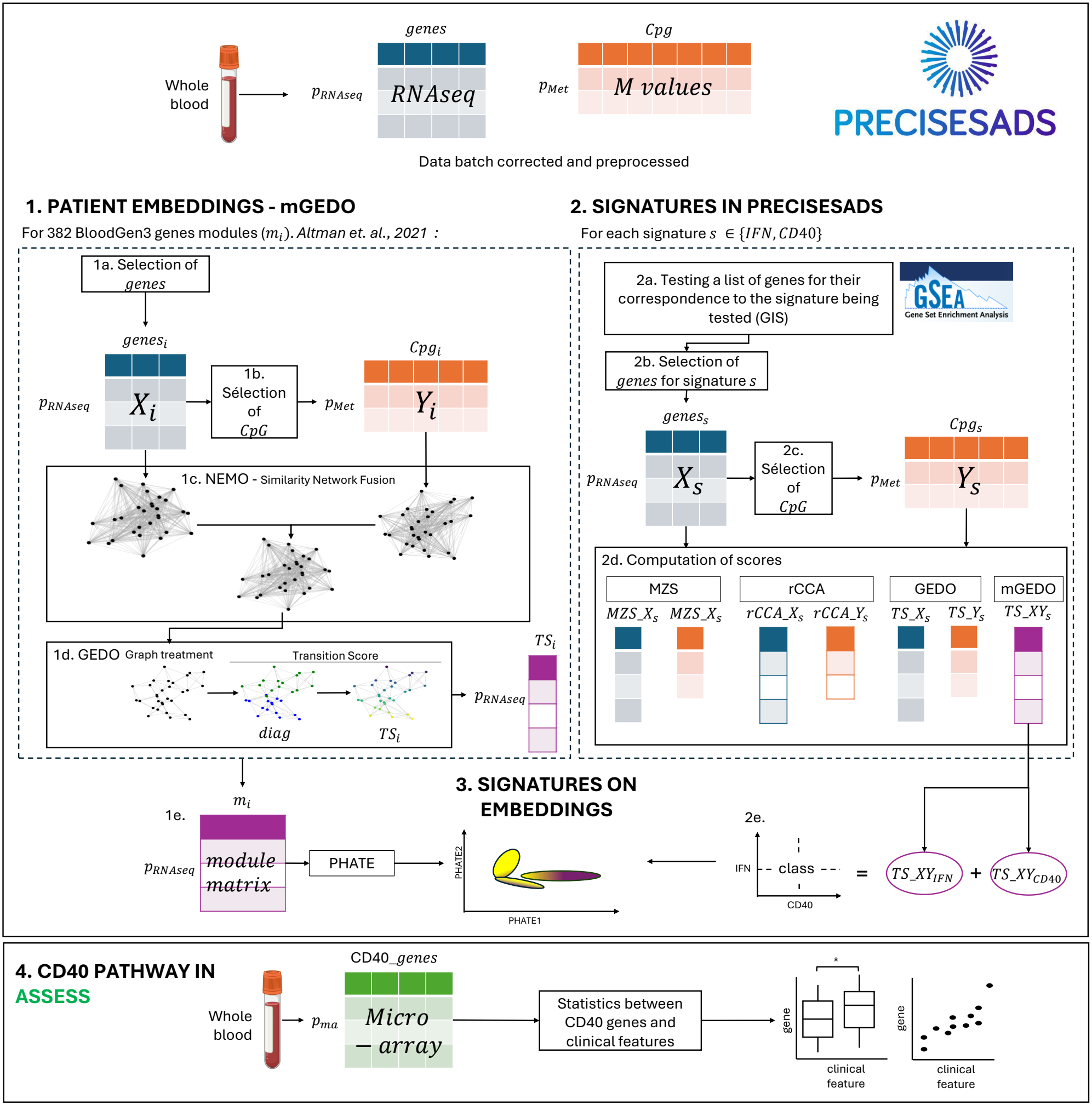
Flowchart of the mGEDO pipeline for the mapping construction and computation of IFN and CD40 signature. Whole-blood RNA-seq and CpG methylation data from PRECISESADS were batch-corrected. **1. MAPPING OF INDIVIDUALS**.For each BloodGen3 gene module (*m*_*i*_): **1a**. The RNA-seq subset *X*_*i*_, corresponding to gene list (*genes*_*i*_), was selected. **1b**. CpGs in regions of interest (*CpG*_*i*_) of *genes*_*i*_ were selected from methylation data (*Y*_*i*_). **1c**. NEMO was used to construct an integrated patient network from RNA-seq (*X*_*i*_) and CpG (*Y*_*i*_) data. **1d**. GEDO was then used to score all patients in this network in a supervised manner by computing each patient’s relative position with respect to CTRL and SjD groups. This produced one Transition Score (*TS* _*i*_) per individual. **1e**. Combining *TS* across all modules (*m*_*i*_) produced a module matrix of 702 individuals (*p*_*RNAseq*_) and 382 gene modules. This matrix was then projected into two dimensions with UMAP and PHATE. **2. SIGNATURES IN PRECISEASDS**. For IFN and CD40 signature (*s*): **2a**. Our automated gene-set selection approach was used to define signature-specific genes of interest (*genes*_*s*_). GIS: Gene Importance Score. **2b**. The RNA-seq subset *X*_*s*_, corresponding to signature genes (*genes*_*s*_), was selected. **2c**. CpGs in regions of interest (*CpG*_*s*_) of signature genes (*genes*_*s*_) were selected from methylation data (*Y*_*s*_). **2d**. Signature scores were computed: (1) Mean z-scores relative to CTRL in *X*_*s*_ (MZS_X_s_) and *Y*_*s*_ (MZS_Y_s_), (2) RNA and CpG components from the rCCA model integrating both *X*_*s*_ and *Y*_*s*_ with the 335 patients with complete data (rCCA_X_s_ and rCCA_Y_s_, respectively), (3) GEDO scoring for each omic separately with Transition Score (TS_X_s_ on *X*_*s*_ and TS_Y_s_ on *Y*_*s*_), (4) Multi-omic scoring with mGEDO, computing a Transition Score accounting for both RNA and CpG data information (TS_XY_s_). **2e**. TS_XY_IFN_ and TS_XY_CD40_ were combined to define CD40-IFN classes. *p*_*RNA*−*seq*_=702, *p*_*Met*_=335. **3. SIGNATURES ON MAPPING**. Signature scores were projected onto PHATE mapping to visualize their enrichment across SjD patients. **4. CD40 PATHWAY IN ASSESS**. CD40 genes in microarray data were crossed with clinical features of the ASSESS cohort, at baseline. *p*_*ma*_=371.

## 3. Results

### 3.1. New two-dimensional mapping reveals a CTRL-to-SjD continuum and SjD subgroups

To get the mapping of CTRL and SjD, first, single omic module matrices were projected in 2 dimensions with PHATE and UMAP. The first was projected from RNA-seq data (Figure 2a,b), and the second was from DNA methylation data (Figure 2c,d). As expected, we observe great heterogeneity within SjD and CTRL, with some cross-category samples. The RNA-seq mapping projected with PHATE (highlighting transitions and branches) shows two branches from CTRL to SjD, highlighting the complex and nonlinear transition structure across patients (Figure 2a).

**Figure 2.**
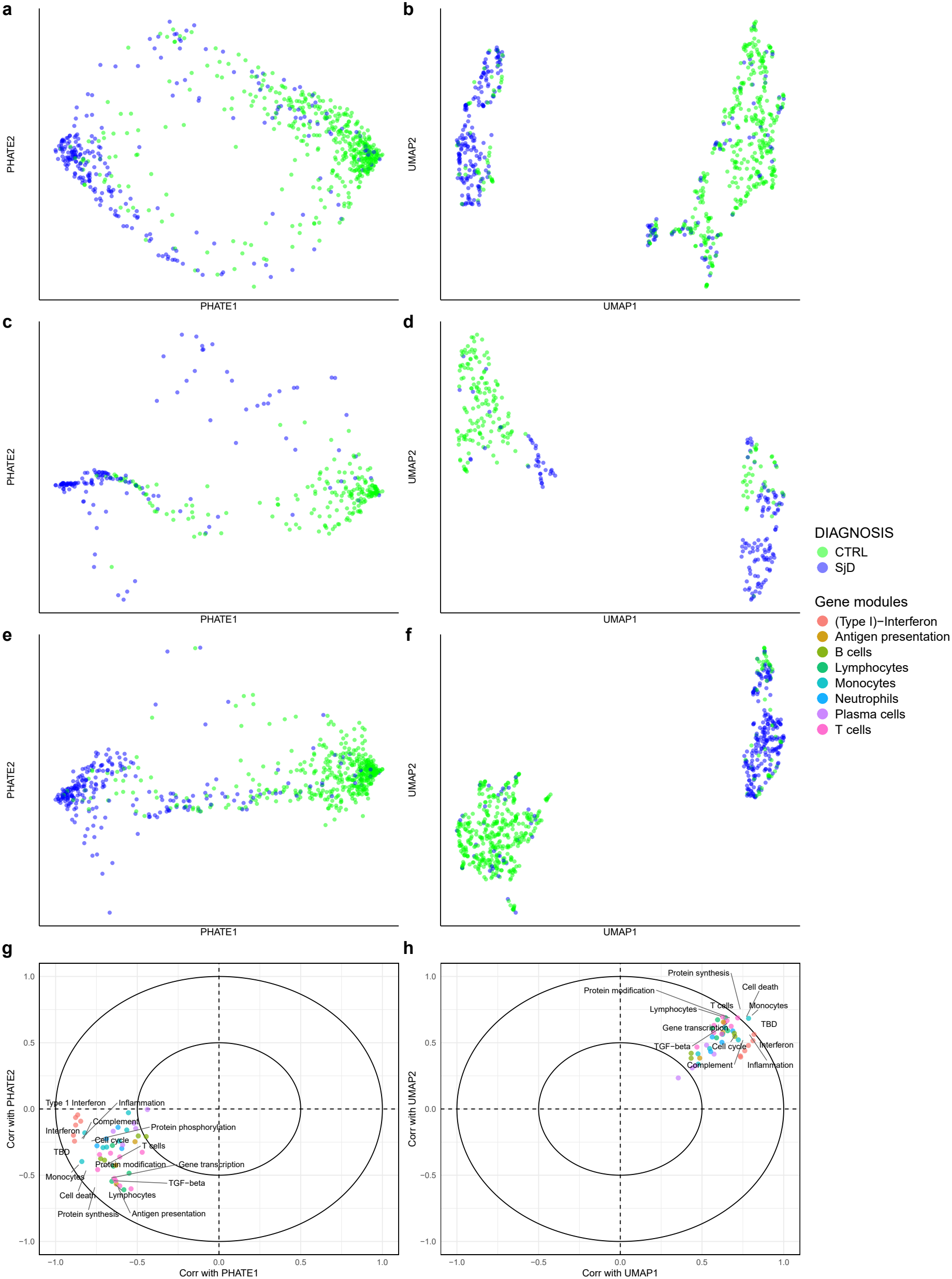
PHATE (left) and UMAP (right) mappings. **a.**and **b**.: projection of patients from the module matrix computed from RNA-seq data with GEDO. **c**. and **d**.: projection of patients from the module matrix computed from CpG methylation data with GEDO. **e**. and **f**.: projection of patients from the module matrix resulting from RNA-seq-CpG integration with mGEDO. **g**. and **h**.: Circos correlation plots representing one point by BloodGen3 gene module related to immunology or immune cell types and its position according to its correlation coefficient with PHATE1 and PHATE2 (left), and UMAP1 and UMAP2 (right) axes. The 50 most important biological functions contributing to mapping construction are annotated (BloodGen3 annotations). TBD: To Be Determined.

With multi-omic integration, integrating information from both gene expression and DNA methylation, transcriptomic data contributed more to the integrated mapping, relative to the methylomic one (Supplementary Notes, Supplementary Figure 1). We also demonstrated that the continuous two-dimensional PHATE mapping obtained from the multi-omic integration better represents the data in comparison to clustering (Supplementary Notes, Supplementary Figure 2). In the multi-omic PHATE mapping, the majority of CTRL individuals cluster together, and SjD patients separate into three principal areas: (1) CTRL-like, really close to the majority of CTRL samples, (2) pure SjD branch, with the majority of SjD patients, far from CTRL, (3) secondary SjD branch (at the bottom-left of Figure 2e), together with some CTRL.

Major biological functions of importance in this separation can be divided into three categories: (1) Interferon (type 1) related pathways, (2) immunological functions, and (3) cell cycle and transcriptional process or protein maturation functions (Figure 2g,h), confirming the biological consistency of the two-dimensional representations.

### 3.2. IFN and CD40 pathway signatures revealed complex gene expressions and CpG methylation alterations

Our new automatic gene selection method allowed us to select genes for:

#### (1) IFN gene set

*IFNA1, IFNB1, IFNA13, IFNA14, IFNA17, IFNA8, IFNA16, IFNA21, IFNA10, IFNA6, IFNA7, IFNA5, IFNA2, IFNAR1, IFNA4, IFNAR2, STAT2, USP18, TYK2, MX1, SOCS1, IRF7, ISG15, IFITM1, JAK1, STAT1*, consistent with previous knowledge on IFN signature (Nezos et al., 2015). 22 out of these 26 genes were actually present in the RNA-seq data (Supplementary Figure 4a,b),

#### (2) CD40 gene set

*CD40, TRAF3, CD40LG, MAP3K14, TRAF2, NFKB2, TRAF6, NFKBIA, TNFAIP3*, all present in RNA-seq data (Supplementary Figure 5a,b).

TS signature method was consistent but managed to separate SjD and CTRL better than MZS and rCCA on the three signatures (Supplementary Notes, Fig. 4d, 5d, 6d).

For the CD40 signature, we tested the contribution of each gene and CpG in the CD40 signature by computing Pearson correlation between each omic feature and the TS_XY_CD40_. The strongest contributor of the *CD40* signature is the *CD40LG* gene (negative correlation < -0.5), and positive contributors are *NFKB2, TNFAIP3, NFKBIA*, and *CD40* with correlation > 0.25. When mixed with CpG, these genes occupied the highest contribution (Supplementary Figure 5e).

Among the nine genes in the CD40 gene set, *CD40, NFKB2, TRAF6*, and *TNFAIP3* expression shows significant over-expression in SjD in comparison to CTRL (p-values = 5.6e-5, 0.003, 0.022, 7.7e-10, respectively). *CD40LG* and *MAP3K14* show significant under-expression in SjD (p-values < 2.22e-16 and 0.006, respectively), with *CD40LG* showing the greatest contrast between SjD and CTRL among all genes tested (Supplementary Figure 6).

As expected, the *CD40* gene correlated strongly with the *TN-FRSF13C* gene, expressing BAFF-receptor (correlation: 0.59, p-value<2.2e-16) but not with BAFF, and *CD40LG* gene was negatively correlated with *TNFSF13B* gene expressing BAFF (correlation: -0.48, p-value<2.2e-16) and poorly with BAFF-receptor, confirming the consistency of our genes with BAFF markers present on B and T cells, respectively.

The three signatures were projected onto the PHATE mapping to test for their co-location among patients. CD40 signature showed a distinct distribution from the IFN signature across individuals. To quantify this difference, the difference between the CD40 and IFN signature was computed, after scaling. We observed both low and high values, indicating IFN- and CD40-predominant individuals. Within the CD40-IFN class, CTRL individuals clustered predominantly as CD40−/IFN−, whereas the mapping displayed a gradual transition from CD40+/IFN −to CD40+/IFN+ and finally CD40− /IFN+, suggesting distinct but partial overlap between CD40- and IFN-predominant patients (Figure 3). Among CD40-related genes, *CD40LG* was the most clearly enriched and representative of the complementarity with the IFN signature in the PHATE mapping, revealing a global expression closely associated with the PHATE2 axis. *CD40* gene showed an expression closer to the IFN signature (Supplementary Figure 8).

**Figure 3.**
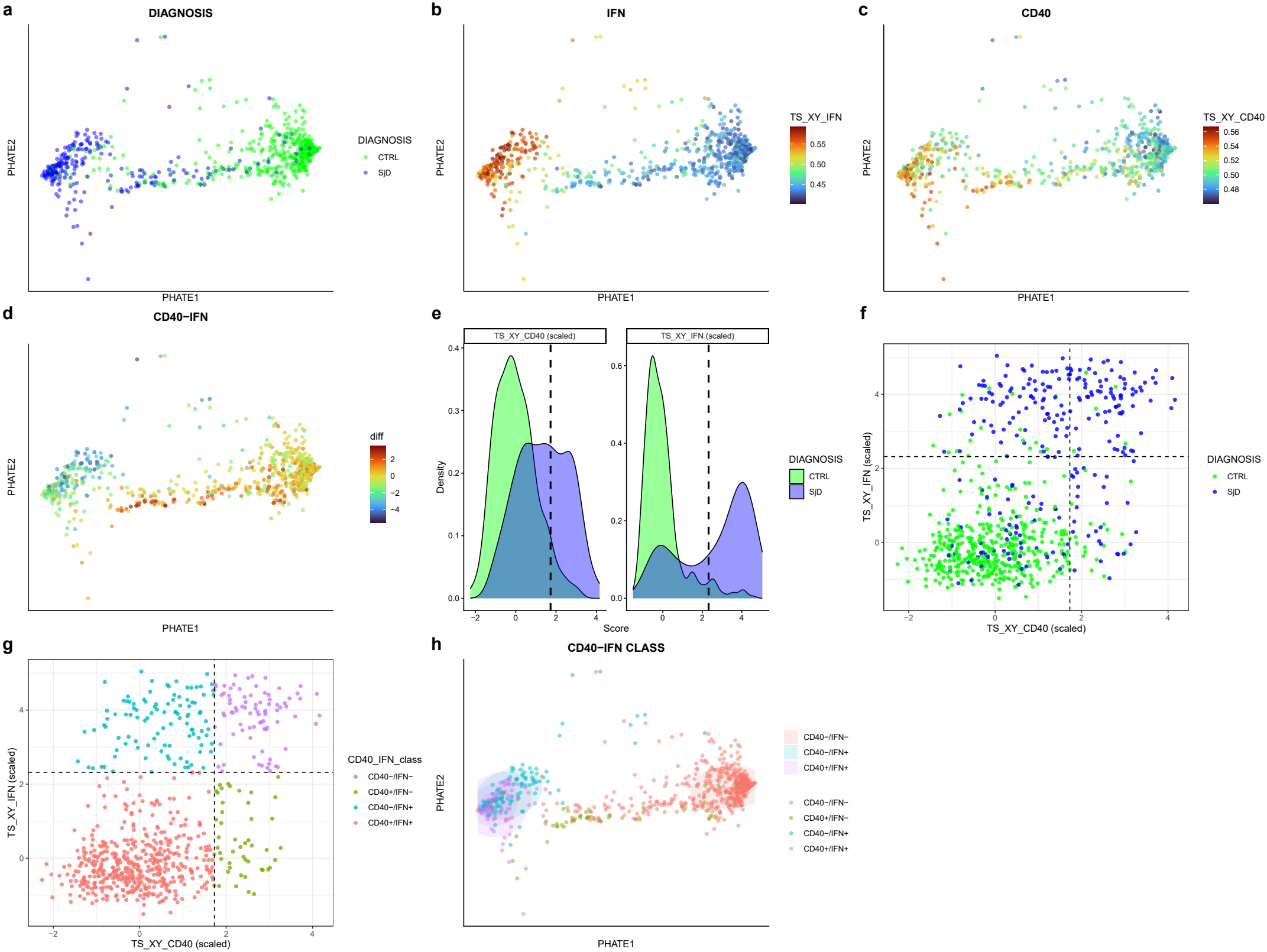
PHATE mapping, colored by: **a.**Diagnosis (SjD or CTRL), **b**. IFN: IFN signature score, **c**. CD40: CD40 signature score, **d**. CD40-IFN signature difference after scaling, **e**. Distribution of TS_XY_CD40_ and TS_XY_IFN_ signature scores, scaled by mean and standard deviation. Black dashed lines: 95th percentile of CTRL individuals (defining positivity for both IFN and CD40 signatures). **f**. IFN signature according to CD40 signature, colored by diagnosis. **g**. IFN signature according to CD40 signature, colored by CD40-IFN class (relative to the positivity of the IFN and CD40 signature). Dark lines: positivity thresholds. **h**. CD40-IFN class (circular areas represent density zones of each class).

### 3.3. IFN and CD40 signatures reveal progressive biological severity in patients with SjD

In the PRECISESADS cohort, among all clinical features tested, five showed significant association with CD40-IFN class after p-value correction: Hypergammaglobulinemia, anti-Ro52 autoantibodies, anti-Ro60 autoantibodies, anti-SSB autoanti-bodies, ESSDAI biological component (corrected p-value < 0.001 in each case). Global ESSDAI shows no association with CD40-IFN class (corrected p-value: 0.743). These five significant features show a very clear progression: from CD40-/IFN-, then CD40+/IFN-, then CD40-/IFN+, and finally CD40+/IFN+ (Figure 4a). In contrast, dryness indicators (dry eyes, dry mouth, focus score, Schirmer test, and salivary flow) were not associated with the CD40-IFN class given the limitation of the low number of observations (Figure 4b, Supplementary Figure 10 and 11, Supplementary Table 2).

**Figure 4.**
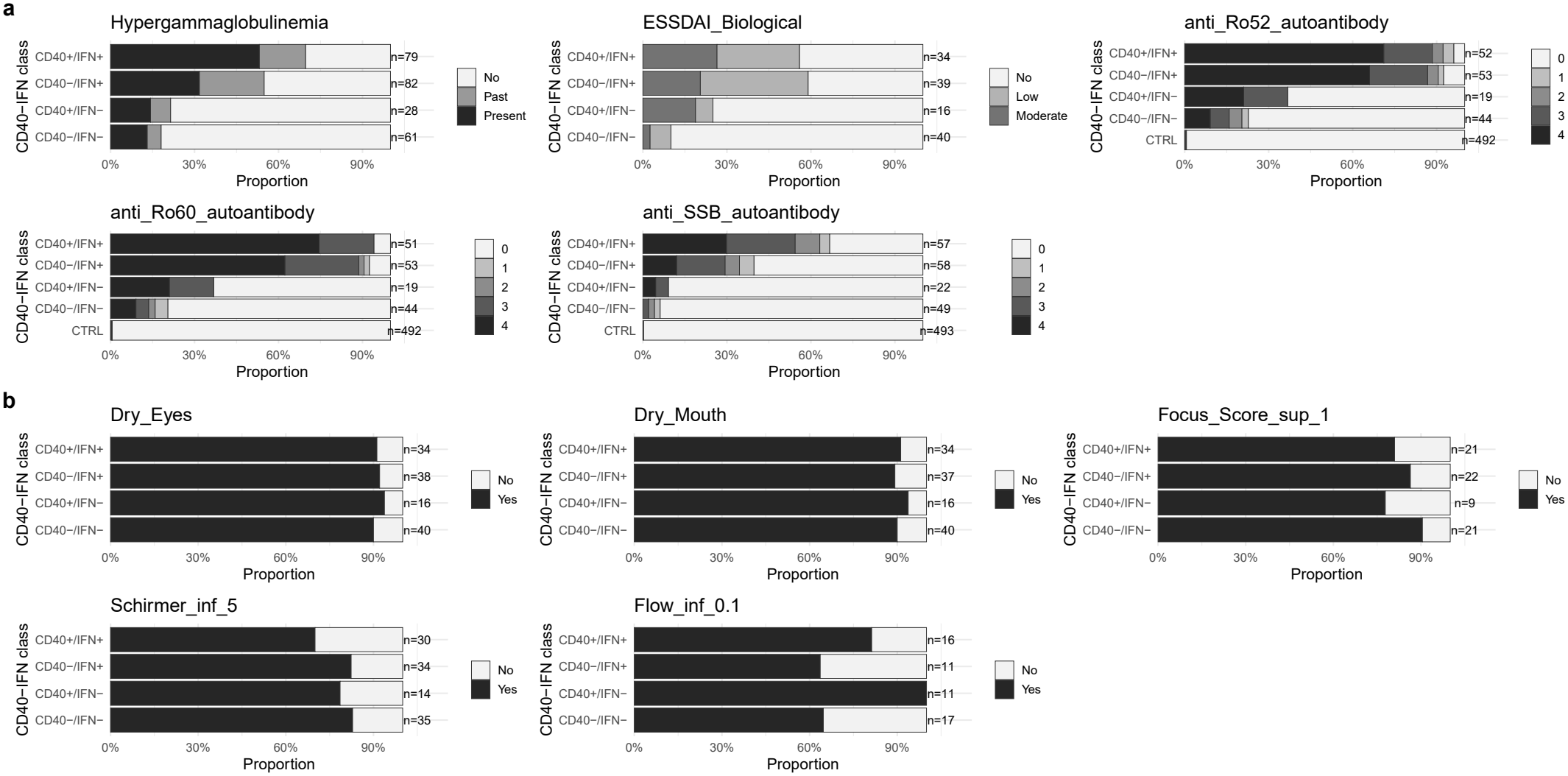
Enrichment of biological and clinical markers across CD40-IFN classes in SjD patients. Proportion: percentage of patients in each class. n: total number of observations in each class. **a.**All variables significantly different across CD40-IFN classes (ESSDAI_Biological: biological component of ESSDAI). **b**. Dryness indicators (Focus_Score_sup_1: Biopsy of the accessory salivary glands with lymphocytic sialadenitis and a focus score greater than or equal to 1, Schirmer_inf_5: Schirmer test less than 5, Flow_inf_0.1: salivary flow less than 0.1 mL/min).

### 3.4. CD40 and CD40LG global expression are associated with cell composition changes

*CD40* and especially *CD40LG* gene expression was associated with several cell population abundances in linear models (See Supplementary Figures 12 and 13 for correlations).

*CD40* expression was positively associated with monocyte concentration (*p* = 7.0 × 10^−4^) and negatively with PMN concentration (*p* = 1.36 × 10^−6^). Within monocytes, *CD40* was positively associated with intermediate monocytes (frequency: *p* = 8.75 × 10^−4^; concentration: *p* = 9.16 × 10^−8^) and negatively with classical monocyte concentration (*p* = 1.66 ×10^−5^). Negative associations were also observed with NKT-like cell concentration (*p* = 0.0168), neutrophil concentration (*p* = 0.0414), and mDC1 frequency (*p* = 0.0.0413). *CD40* was strongly and positively associated with overall B-cell abundance (*p* = 0.00135). *CD40LG* was negatively associated with monocyte (*p* = 7.64 × 10^−15^) and PMN (*p* = 2.95 × 10^−5^) concentrations, especially intermediate monocytes (frequency: *p* = 6.22 × 10^−6^; concentration: *p* = 4.81 × 10^−7^). Within T cells, *CD40LG* expression was positively associated with all major T-cell subset frequencies (all *p* < 5 × 10^−5^), while concentrations showed positive associations with CD4^+^ (*p* < 10^−16^) and CD4^+^CD8^+^ T cells (*p* = 0.0105) and a negative association with CD8^+^ T cells (*p* = 4.12 × 10^−11^). In the B-cell panel, differentiation, CD5×differentiation, and CD5×TACI models were significant.

### 3.5. Association of CD40 signature with clinical and biological manifestations in ASSESS cohort

The associations between *CD40* and *CD40LG* genes with biological activity were confirmed in the ASSESS validation dataset. Even if R-squared values are limited, significant features show that *CD40* gene expression is positively associated with biological systemic activity: IgG, CCL19, Lambda and Kappa chains, gamma globulins, Beta 2 microglobulin, anti-SSA and anti-SSB autoantibodies (see Figure 5a), but not with clinical activity. *CD40LG* gene expression was negatively associated with the same biological features as for CD40, and with CXCL10 (or IP-10), BAFF, and biological and hematological components of ESSDAI at months 24, 48, and 60 after recruitment, despite *CD40LG* correlating positively with absolute lymphocyte count (Figure 5b), but not with clinical activity. Other genes in the CD40 gene set were not associated with dryness indicators: Schirmer test and salivary flow. All statistical results are presented in Supplementary notes.

**Figure 5.**
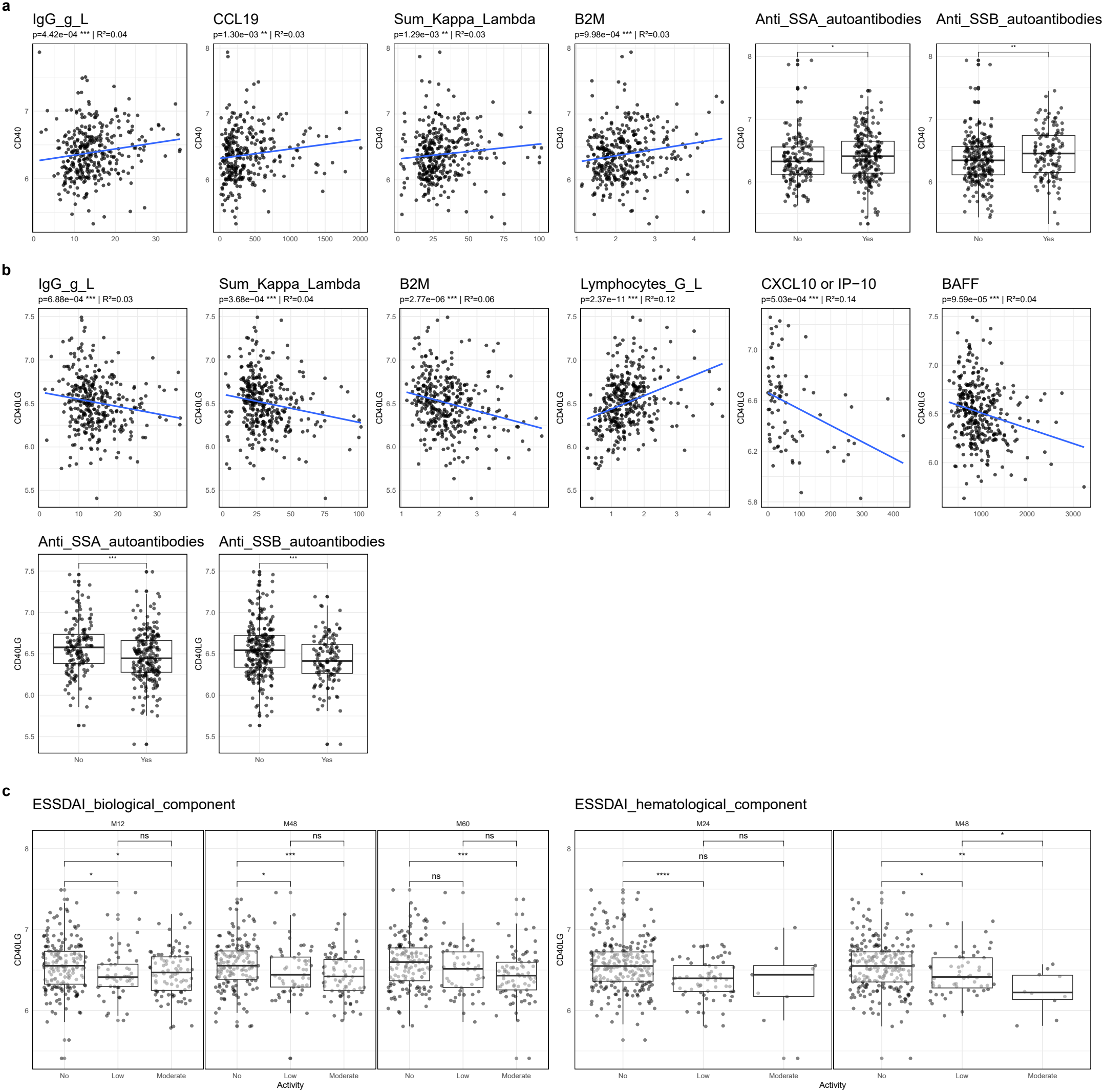
Associations between *CD40* and *CD40LG* gene expression and clinical characteristics -retained from previous systematic statistics-of patients with SjD in the ASSESS cohort. **a.***CD40*, **b**. *CD40LG*, **c**. Association between *CD40LG* expression and ESSDAI biological and hematological components activity. Correlations were tested with Spearman’s test, presented with p-value and proportion of variance explained (r^2^), and differences between classes were tested by the Wilcoxon test. ns: not significant, *: p-value < 0.05, **: p-value < 0.01, ***: p-value < 0.001. B2M: Beta-2 microglobulin; MXX: months after baseline.

## 4. Discussion

Our study provides a multi-omic (transcriptomic and methylomic) characterization of SjD using mGEDO and highlights three main findings: (1) a continuous molecular landscape better captures patient heterogeneity than discrete clustering, (2) DNA methylation substantially contributes to disease stratification and molecular signature construction, and (3) IFN and CD40 signatures represent complementary dimensions of disease mechanisms.

The continuous mGEDO patient landscape allowed us to observe marked heterogeneity among both SjD patients and controls, including CTRL-like SjD patients fulfilling diagnostic criteria despite near-normal molecular profile. Beyond these biological insights, this work demonstrates the ability of mGEDO and the supervised Transition Score to quantify disease-related molecular alterations into a continuous pathological axis.

The integration of DNA methylation was motivated by previous works (Imgenberg-Kreuz et al., 2016; Teruel et al., 2021). Our analyses confirm that methylation contributes substantially to patient stratification and pathway signatures, with some of the CpG sites accounting positively or negatively in signature scores.

Consistent with extensive literature, type I IFN remained the dominant molecular axis separating SjD from controls and was associated with the strongest immunological abnormalities, including BAFF activation, B-cell hyperactivity, autoantibody production, and increased disease activity (Brkic et al., 2013; Maria et al., 2014). However, IFN alone did not involve all patients (Teruel et al., 2021).

Rather than being redundant, IFN and CD40 signatures appear complementary, especially in IFN-negative patients. IFN acts upstream by promoting BAFF production, B-cell activation and Tfh differentiation, whereas CD40 reflects active T–B cell interactions required for germinal center formation, plasma-cell differentiation and ectopic lymphoid structures (Crotty, 2011; Nocturne and Mariette, 2018; Boyman and Sprent, 2012). Accordingly, CD40+/IFN+ patients displayed the strongest biological abnormalities, including hypergamma-globulinemia, autoantibody positivity, and increased biological ESSDAI domains, suggesting that CD40–IFN classes were more strongly associated with biological markers than with patient-reported symptoms, consistent with previous observations (Bodewes et al., 2018; Teruel et al., 2021; Del Papa et al., 2021). IFN+/CD40– patients may represent an earlier stage of disease before sustained adaptive immune activation, whereas CD40+/IFN– profiles may reflect tissue-driven immune activation within salivary glands independently of systemic IFN activation (Psarras et al., 2017; Tang et al., 2024; Rivière et al., 2020). Genetic variants of BAFF are also responsible for a higher risk of SjD, independently of the IFN pathway (Beydon et al., 2026).

We described a CD40 signature which co-located with the expression of BAFF, suggesting a link with B-cell activation or survival, or associated cell composition changes. *CD40* gene expression increased along with BAFF-receptor one, which could be explained by B cells proportion changes in circular blood. *CD40* gene expression correlated with multiple markers of disease activity, including immunoglobulins, free light chains, β_2_-microglobulin, gamma globulins, SSA/SSB autoantibodies, CCL19 (which contributes to tertiary lymphoid structure organization and chronic glandular inflammation (Liu et al., 2019)) and biological ESSDAI domains, consistent with enhanced B-cell and myeloid activation in SjD (Gottenberg et al., 2013; Liu et al., 2019; Rivière et al., 2020).

In contrast, *CD40LG* expression decreased in peripheral blood of SjD and was negatively correlated with both expression of the BAFF gene and the level of BAFF protein in blood. One might hypothesize that this reflects the balance between T cells (expressing CD40LG) and myeloid cells (expressing BAFF) in peripheral blood. It cannot be concluded that these observations are related to actual gene expression programs or are simply a reflection of changes in cellular composition, as the *CD40LG* gene correlated positively with lymphocyte counts and negatively with several biological disease activity markers, including IP-10. *CD40LG* is primarily expressed by activated CD4^+^ T cells and Tfh2 cells (Liu et al., 2022), which are redistributed to salivary glands where they participate in ectopic lymphoid structure formation and interact with CD40-expressing B cells (Rivière et al., 2020; Verstappen et al., 2021). Although circulating soluble CD40L concentrations are elevated in SjD blood (Goules et al., 2006), the reduction of *CD40LG* transcripts in the blood may reflect the migration of activated T-cell populations from the circulation to inflamed tissues, especially in some SjD subgroups characterized by T-lymphopenia (Soret et al., 2021), consistent with our observed results (positive association with CD4+T cells). To reinforce this hypothesis, *CD40LG* expression was strongly associated with frequencies of T cells -CD4+/CD8+ balance in particular-in flow cytometry experiments, suggesting the link with T cell orientation and migration. Some authors described also a down regulation of *CD40LG* expression in some T helper cells after CD40-CD40L contact, which can explain partially the down expression of that gene in SjD patients (Lee et al., 2002).

Together, these observations support IFN and CD40 signatures as distinct but interconnected dimensions of immune activation in SjD.

Several limitations should be acknowledged. First, published gene sets used for validation may not be fully independent of the gene-selection procedure. Second, whole-blood transcriptomic and methylomic profiles cannot distinguish cell-type-specific from compositional effects (Houseman et al., 2012; Titus et al., 2017). In general, the proposed signatures remain to be validated experimentally.

Future studies will focus on *CD40LG* and *CD40LG* expression to determine whether this expression is entirely due to changes in cellular composition, or whether actual transcriptional regulatory programs are at play. Extending this strategy to additional pathways may further refine molecular stratification and improve our understanding of SjD pathophysiology. From a clinical perspective, the CD40–IFN framework may prove useful for clinical severity stratification and for studying lymphoma risk over time. These results could be replicated in clinical trials to test for differential treatment efficacy, as proposed by Chevet et.al. (Chevet et al., 2025), and support new discoveries in precision medicine of SjD.

## Acknowledgements and affiliations

This work was funded by Bio Logbook. The French Ministry of Research has awarded this work a grant under the program CIFRE with the reference 2023/0522. The present CIFRE program supports a thesis and involves an academic partner (University of Western Brittany), an industrial partner (Bio Logbook, 2018, www.biologbook.fr), which is developing solutions for precision healthcare, and CB, the PhD student. This work was supported by the Innovative Medicines Initiative Joint Undertaking under Grant Agreement Number 115565, the resources of which are composed of financial contributions from the European Union’s Seventh Framework Program (FP7/2007–2013) and EFPIA companies in kind.

RB is the founder of Bio Logbook, and this had no impact in the design, the data collection, the analysis, the conclusions, the decision to publish, or the preparation of the manuscript. All other authors have no conflict of interest.

This work was also presented at the 38th Congress of the French Society of Rheumatology and published in *Revue du Rhumatisme* (https://doi.org/10.1016/j.rhum.2025.10.460) in 2025, at the International Sjögren’s Symposium and to the American College of Rheuymatology congress in 2026 (https://acrabstracts.org/abstract/type-i-interferons-and-cd40-complementary-pathways-revealed-by-mgedo-in-distinct-transcripto-methylomic-profiles-of-patients-with-sjogrens-disease-in-the-precisesads-cohort/).

## Key messages

### What is already known on this topic

- Sjögren’s disease is molecularly heterogeneous, and interferon signaling is a major but incomplete explanation of disease mechanisms.
- Conventional clustering often yields unstable or non-concordant patient subgroups across studies.
- CD40 pathway is a biologically relevant therapeutic candidate and could help for refining patient stratification beyond interferon status alone.

### What this study adds

- mGEDO maps SjD and CTRL along a continuous two-dimensional molecular landscape rather than into discrete clusters.
- IFN remains the dominant axis of disease severity, but CD40 provides complementary information, especially in IFN-negative patients.
- Combined IFN/CD40 stratification highlights a biological severity gradient, with double-positive patients showing the strongest immune dysregulation.
- IFN and CD40 signature could help distinguish partially uncoupled innate and adaptive immune programs.
- The lower expression of *CD40LG* in whole blood may reflect T cell migration to salivary or lacrimal glands and could be a marker of disease activity.

### How this study might affect research, practice or policy

- Multi-omic continuous representations may improve patient stratification for future precision-medicine trials in SjD or other diseases.
- mGEDO seems to be a powerful framework for multi-omic integration for global patient analysis and the study of signalling pathways.
- IFN and CD40 signatures could help to stratify patients with SjD at a molecular and clinical level.
- This framework may support more targeted therapeutic strategies and longitudinal studies, including lymphoma-risk stratification.

## Research data

The PRECISESADS data underlying this article will be shared on reasonable request to the corresponding author. All data included in our study are available upon request at ELIXIR Luxembourg, with the permanent link: doi:10.17881/th9v-xt85. Additionally, the relationship between gene expression and DNA methylation datasets can be explored at http://bioinfo.genyo.es/precisesadsdata/. For ASSESS cohort, anonymized data, along with the data dictionary, will be available upon publication. All access requests must be directed to Xavier Mariette. Data transfer will be subject to an access protocol and will require authorization from the scientific board of the ASSESS cohort. Data sharing will be contingent upon a data sharing agreement before any transfer, to ensure all users of the data adhere to the legal requirements of using personal data.

## Code availability

The source code to run mGEDO and reproduce figures is available under the GNU General Public License v3.0 at https://github.com/biologbook/GEDO.

